# Reparameterization of PAM50 expression identifies novel breast tumor dimensions and leads to discovery of a breast cancer susceptibility locus at 12q15

**DOI:** 10.1101/133397

**Authors:** Michael J Madsen, Stacey Knight, Carol Sweeney, Rachel Factor, Mohamed Salama, Venkatesh Rajamanickam, Bryan E Welm, Sasi Arunachalam, Brandt Jones, Kerry Rowe, Melissa Cessna, Alun Thomas, Lawrence H Kushi, Bette J Caan, Philip S Bernard, Nicola J Camp

## Abstract

It is well-known that breast tumors exhibit different expression patterns that can be used to assign intrinsic subtypes – the PAM50 assay, for example, categorizes tumors into: Luminal A, Luminal B, HER2-enriched and Basal-like – yet tumors are often more complex than categorization can describe. We used 911 sporadic breast tumors to reparameterize expression from the PAM50 genes to five orthogonal tumor dimensions using principal components (PC). Three dimensions captured intrinsic subtype, two dimensions were novel, and all replicated in 945 TCGA tumors. By definition dimensions are independent, an important attribute for inclusion in downstream studies exploring effects of tumor diversity. One application where tumor subtyping has failed to provide impact is susceptibility genetics. Germline genetic heterogeneity reduces power for gene-finding. The identification of heritable tumor characteristics has potential to increase homogeneity. We compared 238 breast tumors from high-risk pedigrees not attributable to *BRCA1* or *BRCA2* to 911 sporadic breast tumors. Two PC dimensions were significantly enriched in the pedigrees (intrinsic subtypes were not). We performed proof-of-concept gene-mapping in one enriched pedigree and identified a 0.5 Mb genomewide significant region at 12q15 that segregated to the 8 breast cancer cases with the most extreme PC tumors through 32 meioses (p=2.6×10^−8^). In conclusion, our study: suggests a new approach to describe tumor diversity; supports the hypothesis that tumor characteristics are heritable providing new avenues for germline studies; and proposes a new breast cancer locus. Reparameterization of expression patterns may similarly inform other studies attempting to model the effects of tumor heterogeneity.

## Introduction

The discovery of distinctive gene expression patterns(1) and breast tumor intrinsic subtypes (Luminal A, Luminal B, HER2-enriched, and Basal-like) have illustrated different paths to tumorigenesis and associations with clinical endpoints(2,3). These landmark studies underscored expression as an important attribute of a tumor, with clinical relevance. However, categorization of tumors in to mutually excusive intrinsic subtype based on similarity to archetypal tumors may be an oversimplistic use of these gene expression patterns. Profiles often exhibit various aspects that resemble different archetypal subtypes, and this admixture is lost in the current end-use of gene features. Here we revisit how to present expression data from selected features (i.e. genes previously defined as classifiers) for more flexible end-uses of these important discriminators. Improved extraction of expression variation from feature sets optimizes the potential to advance our understanding of tumor diversity and its usage across diverse domains, such as gene mapping, prediction and treatment.

The high-risk pedigree design has been instrumental in the mapping and discovery of germline susceptibility genes for breast cancer(4,5). Critical to success is an informative phenotype. Power is optimized for phenotypes where the underlying genetics are homogeneous. A focus on early onset disease led to evidence for the high penetrance genes *BRCA1* and BRCA2(6,7). However, beyond these early successes, little progress has been made with pedigree-based gene-mapping for pedigrees not attributable to *BRCA1* or *BRCA2* (non-*BRCA1/2*); genetic heterogeneity remains a major obstacle. A ‘same-gene-same-molecular-subtype’ hypothesis is supported by the fact that *BRCA1* tumors have distinct expression profiles(8) and are most often Basal-like(9). Further, small *non-BRCA1/2* family studies (18 tumors in 8 families(10), and 23 tumors in 11 families(11)) observed some tumor subtype patterning consistent with an ability to partition non-*BRCA1/2* tumors. However, definitive evidence for this hypothesis remain to be defined. To our knowledge, here we present the largest tumor study in *non-BRCA1/2* families (238 tumors in 11 pedigrees). We use the PAM50(12) for gene expression and explore whether categorical intrinsic subtypes or reparameterized tumor expression dimensions are enriched in pedigrees. Under the ‘same-gene-same-molecular-subtype’ hypothesis such tumor phenotypes would reflect inherited susceptibility and be powerful for gene mapping.

## Materials and Methods

### Population-based tumors: the LACE and Pathways (LACE/Pathways) Studies

The Life After Cancer Epidemiology (LACE) and Pathways Studies are prospective cohort studies of breast cancer prognosis(13, 14) Briefly, in the LACE Study, women were enrolled at least 6 months after diagnosis with Stage I-IIIb breast cancer, with baseline data collection in 2000. In the Pathways Study, 4,505 women at all stages of breast cancer were enrolled between 2006 and 2013, on average about two months after diagnosis. In these studies, most or all women were diagnosed with breast cancer in the Kaiser Permanente Northern California healthcare system; a small proportion of women in the LACE Study were also enrolled in the state of Utah. Both cohorts were sampled as broadly representative of breast cancers in the general population and participants were enrolled without regard to family history of cancer. A stratified random sample from the combined LACE/Pathways cohorts was selected for acquisition of primary tumor punches from FFPE blocks(15). Common hormone receptor positive, HER2-negative breast cancers, by immunohistochemistry subtypes, were sampled at a lower frequency. In this study, expression data for tumors from the 911 Caucasian women was used (selected to match the ethnicity in the Utah pedigrees). Survey weights were provided to address the decreased sampling rate of certain immunohistochemistry subtypes.

Gene expression data was generated using the PAM50 RT-qPCR research assay(12) in the Bernard Lab at the Huntsman Cancer Institute(15). For each tumor sample a calibrated log-expression ratio was produced for each gene, producing an expression matrix. Centroid-based algorithms were used to generate quantitative normalized subtype scores for each of the four clinical subtypes plus a subtype characteristic of normal tissue (“Normal-like”). These subtype scores represent the correlation with “prototypic” breast tumors for each of the subtypes. As per standard protocol, each tumor was categorized according to its maximal subtype score. Each tumor was additionally assigned quantitative proliferation, progesterone receptor (PGR), estrogen receptor (ESR) and ERRB2 expression scores(16).

### Derivation of expression dimensions using Principal Component (PC) Analysis

To explore alternate representation of tumor expression in the PAM50 genes, we used principal component (PC) analysis to identify tumor expression dimensions that explained the majority of variance. Principal components analysis was performed on the PAM50 expression matrix of the 911 population tumors using core packages of R version 3.1.1. To address the stratified sampling of the LACE/Pathways tumors, a sampling strategy incorporating survey weights was used. A weighted random sample with replacement to the correct size (N=911) was performed prior to PC analysis. This weighting procedure was repeated 10,000 times and the resulting PCs from each iteration were aligned as necessary, then averaged and centered. Each PC is a linear combination of the gene expression across the 50 genes, and can be utilized as a quantitative variable. Stacked intrinsic-subtype-specific histograms were used to explore patterns between each quantitative PC and the categorical intrinsic subtypes. Kendall’s tau coefficient was used to quantify the correlation of each PC to the PAM50 quantitative scores for proliferation, PGR, ESR and ERBB2 expression.

### Replication in the The Cancer Genome Atlas (TCGA) data

The TCGA breast tumor expression data was used as a replication study for the established dimensions in the LACE/Pathways data. We used RNA sequencing data for 745 breast tumors from Caucasian women in the TCGA Breast Invasive Carcinoma project. Standardized FPKM values for RNA sequencing transcriptome data and intrinsic subtype were downloaded from the National Cancer Institute GDC portal. The PC rotation matrix derived from the LACE/Pathways data was applied to log-transformed standardized FPKM expression values to establish the defined PC dimensions in the TCGA data. Similarly, stacked intrinsic-subtype-specific histograms were generated. Further, we investigated the possibility that novel PC dimensions were representative of single genes elsewhere in the genome using comparisons of PC variables and individual genes across the genome using Pearson product moment (*ρ*).

### High-risk breast cancer pedigrees: Identification, selection and acquisition of materials

High-risk breast cancer pedigrees were identified in the Utah Population Database (UPDB) through record linkage of a 16-generation genealogy and statewide cancer records from the Utah Cancer Registry (UCR). High-risk pedigrees were defined based on a statistical excess of breast cancer (p<0.05; example Figure 1a). Pedigrees known to be attributable to *BRCA1/2* from previous Utah studies (screen positive or linked to chromosomes 17q21 or 13q13) were removed from consideration. Pedigrees with fewer than 15 meioses between cases were also removed as these lack power for gene-finding(17). Record linkage between the UPDB and electronic medical records in the University of Utah and Intermountain Healthcare systems allowed identification of tumor blocks. Twenty-five *non-BRCA1/2* extended high-risk pedigrees were identified, each with a minimum of 15 available tumors (Table 1). Matched tumor and grossly uninvolved (GU) formalin-fixed paraffin-embedded (FFPE) breast tissue blocks were retrieved for pathological review and acquisition of tumor punches (ideal: 4×1.5 mm tumor punches; minimum 1 punch) and GU scrolls (ideal: 7×15 μm full face GU scrolls; minimum 4 scrolls) was performed. This resulted in 391 quality-controlled paired tumor-GU tissue samples obtained from the Intermountain BioRepository (N=354) and the University of Utah Department of Pathology (N=37). In parallel, living breast cancer cases within the 25 high-risk pedigrees were invited to participate, including a blood draw. Eleven high-risk pedigrees contained the vast majority of the tissue samples (N=245). These 11 most informative pedigrees were the focus of this study. Nucleic acid extraction was performed as described previously(16). After quality control, 238 breast cancer cases had both quality controlled tumor RNA and germline DNA available (45 blood-derived, 1 from saliva, the remaining from GU breast tissue). All women were Caucasian. Ethical approvals for the study were governed by IRBs at the University of Utah and Intermountain Healthcare.

**Figure 1.**
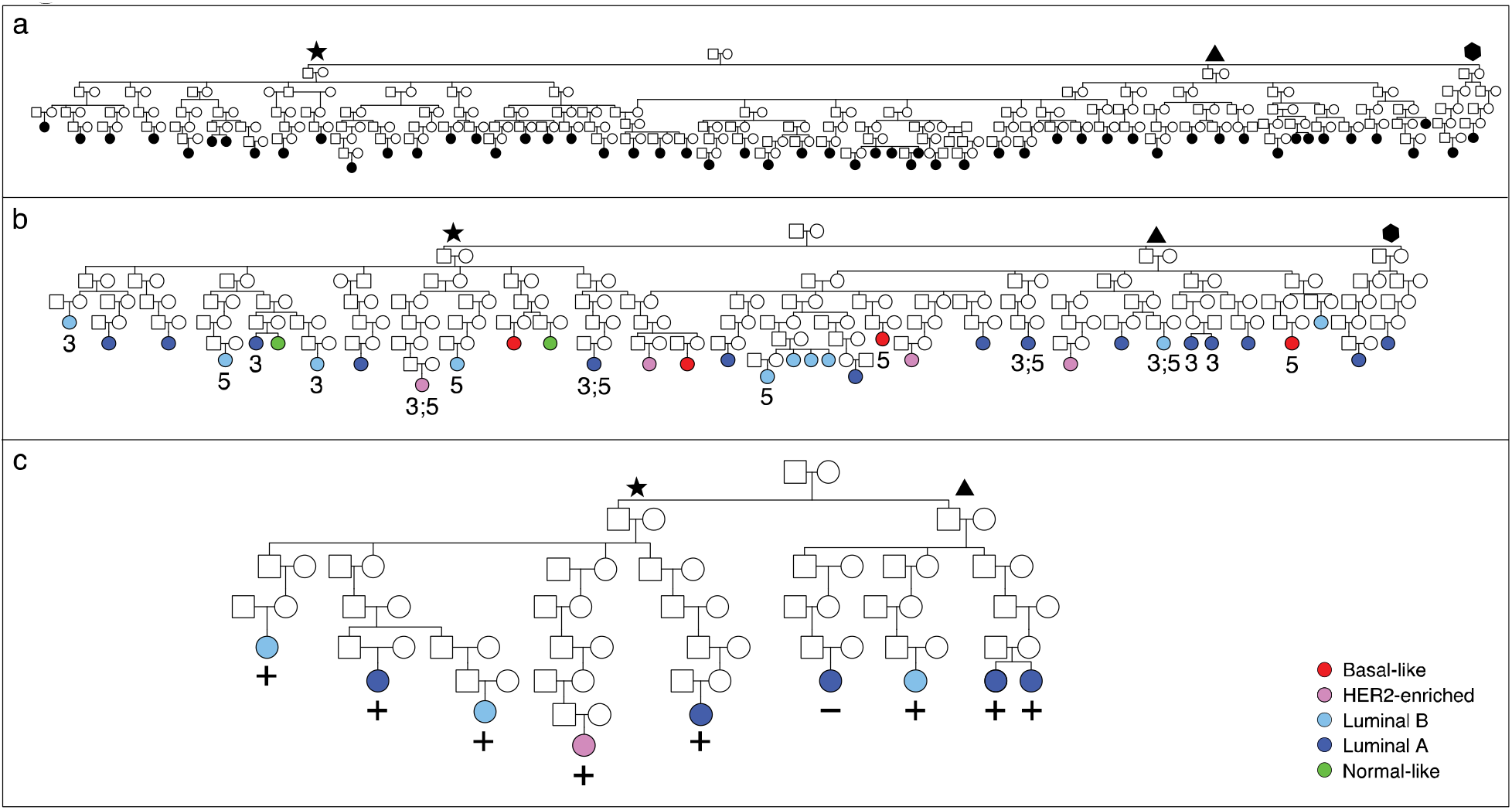
Example Utah high-risk breast cancer pedigree 1817. **a.** Confirmed and sampled breast cancer cases are indicated in black (55 sampled out of 138 total confirmed UCR cases). Star, triangle and hexagon symbols indicate pedigree branches. **b.** shows only those cases from (a) with tumor expression data available and indicates PAM50 intrinsic subtype by color. Cases whose tumors are extreme for PC3 are indicated by ‘3’; extreme for PC5 are indicated by ‘5’. **c.** shows only the PC3-extreme cases from (b). A ‘+’ indicates those cases that share the genomewide significant region at 12q15.

**Table 1:**
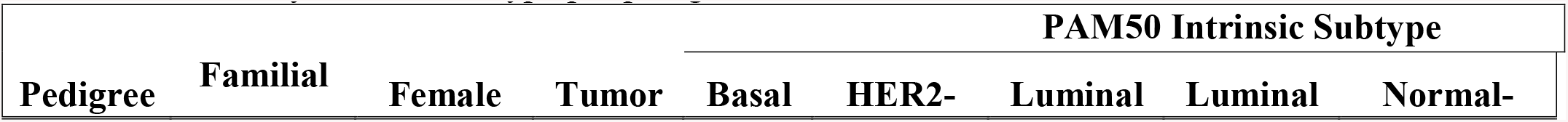
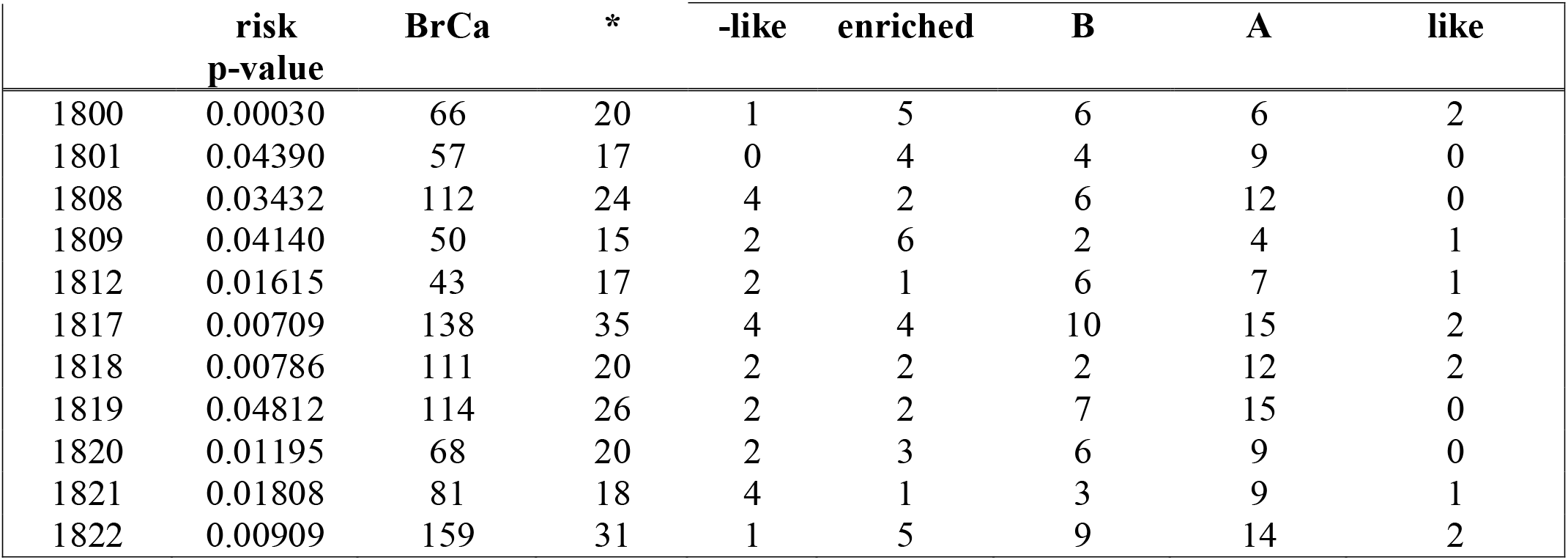
Summary of the 11 high-risk Utah pedigrees. Tumor count by intrinsic subtype per pedigree.

| Pedigree | Familial | Female | Tumor | PAM50 Intrinsic Subtype |  |  |  |  |
| --- | --- | --- | --- | --- | --- | --- | --- | --- |
|  |  |  |  | Basal | HER2- | Luminal | Luminal | Normal- |

|  | risk<br>p-value | BrCa | * | -like | enriched | B | A | like |
| --- | --- | --- | --- | --- | --- | --- | --- | --- |
| 1800 | 0.00030 | 66 | 20 | 1 | 5 | 6 | 6 | 2 |
| 1801 | 0.04390 | 57 | 17 | 0 | 4 | 4 | 9 | 0 |
| 1808 | 0.03432 | 112 | 24 | 4 | 2 | 6 | 12 | 0 |
| 1809 | 0.04140 | 50 | 15 | 2 | 6 | 2 | 4 | 1 |
| 1812 | 0.01615 | 43 | 17 | 2 | 1 | 6 | 7 | 1 |
| 1817 | 0.00709 | 138 | 35 | 4 | 4 | 10 | 15 | 2 |
| 1818 | 0.00786 | 111 | 20 | 2 | 2 | 2 | 12 | 2 |
| 1819 | 0.04812 | 114 | 26 | 2 | 2 | 7 | 15 | 0 |
| 1820 | 0.01195 | 68 | 20 | 2 | 3 | 6 | 9 | 0 |
| 1821 | 0.01808 | 81 | 18 | 4 | 1 | 3 | 9 | 1 |
| 1822 | 0.00909 | 159 | 31 | 1 | 5 | 9 | 14 | 2 |

*\*Tumor samples with PAM50 data that passed QC. Three tumors belonged to two pedigrees and one tumor belonged to three pedigrees.*

### PAM50 gene expression in high-risk pedigrees and comparison to population tumors

Parallel to the tumors from the LACE/Pathways studies, expression data for the 238 tumors in the 11 *non-BRCA1/2* high-risk pedigrees was also generated using the PAM50 RT-qPCR research assay in the Bernard Lab at the Huntsman Cancer Institute (example of Figure 1b).

Intrinsic subtype and proliferation, PGR, ESR and ERRB2 expression scores were assigned as per standard protocol. Principal component variables, as defined in the LACE/Pathways population data, were used to generate PC scores (dimensions) for high-risk pedigree tumors.

Under the null hypothesis that germline genetics do not influence tumor expression, tumors in the pedigrees are independent and the patterns observed should match distributions from the general population. Rejection of the null is consistent with a role for germline variants in tumor expression, and provides potential heritable expression phenotypes. The distribution of intrinsic subtypes for the LACE/Pathways data, corrected for the stratified sampling criteria, has previously been determined(15). Differences in the distribution of intrinsic subtypes between pedigree and LACE/Pathways tumors was evaluated using a chi-squared goodness of fit test. Differences in quantitative PC scores between pedigree and population tumors was determined using a weighted t-test to account for the LACE/Pathways sampling weights. For bi-modal PCs a likelihood ratio test of proportions was implemented. Tests were repeated for each of the 11 pedigrees separately, with a Bonferroni correction applied to account multiple testing (*α*=0.0045).

### Proof-of-concept: Shared Genomic Segment (SGS) with novel PC dimensions

We performed SGS gene mapping in one Utah high-risk pedigree as a case-study to explore the utility of potential novel heritable dimensions. The pedigree was selected based on harboring tumors with PC dimensions significantly different than that expected. SGS analysis requires high-density single nucleotide polymorphism (SNP) data. We used the OmniExpress high-density SNP array with genoptyes called using standard Illumina protocols. SNP quality control included: duplicate check, sex check, SNP call-rate (95%), sample call rate (90%, more liberal due to the FFPE DNA), and failure of Hardy-Weinberg equilibrium (p≤1×10^−5^). SGS analysis identifies statistically significant chromosomal regions shared by multiple, distant relatives. It is based on evaluating identity-by-state (IBS) sharing at consecutive SNP loci, with segregation from a common ancestor implied if the observed sharing is significantly longer than expected by chance(17,18). The method was developed specifically for extended pedigrees, with power gained from the unlikely event that long segments are inherited across a large number of meioses by chance(17). Statistical significance for an SGS chromosomal region is determined empirically using a gene-drop approach. Pairs of haplotypes are randomly assigned to pedigree founders according to the haplotype distribution. Mendelian segregation and recombination are simulated to generate genotypes for all pedigree members. 1000Genomes Project(19) genoptye data were used to estimate a graphical model for linkage disequilibrium (LD)( 20), providing a probability distribution of chromosome-wide haplotypes in the population. The Rutgers genetic map(21) was used for a genetic map for recombination, with interpolation based on physical base pair position for SNPs not represented. Once the gene-drop is complete, simulated SNP genotypes for the individuals of interest are used to determine chance sharing. The gene-drop procedure is repeated tens of millions of times to estimate the significance of the observed sharing. We considered tumors to be “PC-extreme” if they were above the 90^th^ population percentile. We performed genomewide SGS analyses iteratively on ordered subsets of breast cancer cases, beginning with those with the two most extreme PC values, expanding one case at a time, and stopping when all PC-extreme tumors had been considered. Genomewide significance thresholds for SGS that account for multiple testing (all subsets and all chromosomes) have been described previously(22). Briefly, −log_10_ p-values for all chromosomes and across all subsets follows a gamma distribution, with parameters that vary based on the number of cases and the structure of a pedigree. Based on the assumption that the vast majority of all sharing in a genome is under the null, parameters for the gamma distribution can be estimated from the real data and pedigree structure. Pedigree-specific genomewide significance thresholds are then derived from the appropriate gamma distribution using the theory of large deviations.

## Results

### Novel breast tumor dimensions

From the 911 population tumors, PC analysis identified five PCs that accounted for 68% of the total expression variance (30.5%, 18.9%, 10.2%, 5.3%, 3.2% explained by PCs 1–5 respectively). Components beyond PC5 explained diminishing amounts of variance. The relationship between PCs 1–5 and common intrinsic subtypes is illustrated in Figure 2 and gene coefficients (eigenvectors) are shown in Supplemental Table 1 (S1 Table).

**Figure 2.**
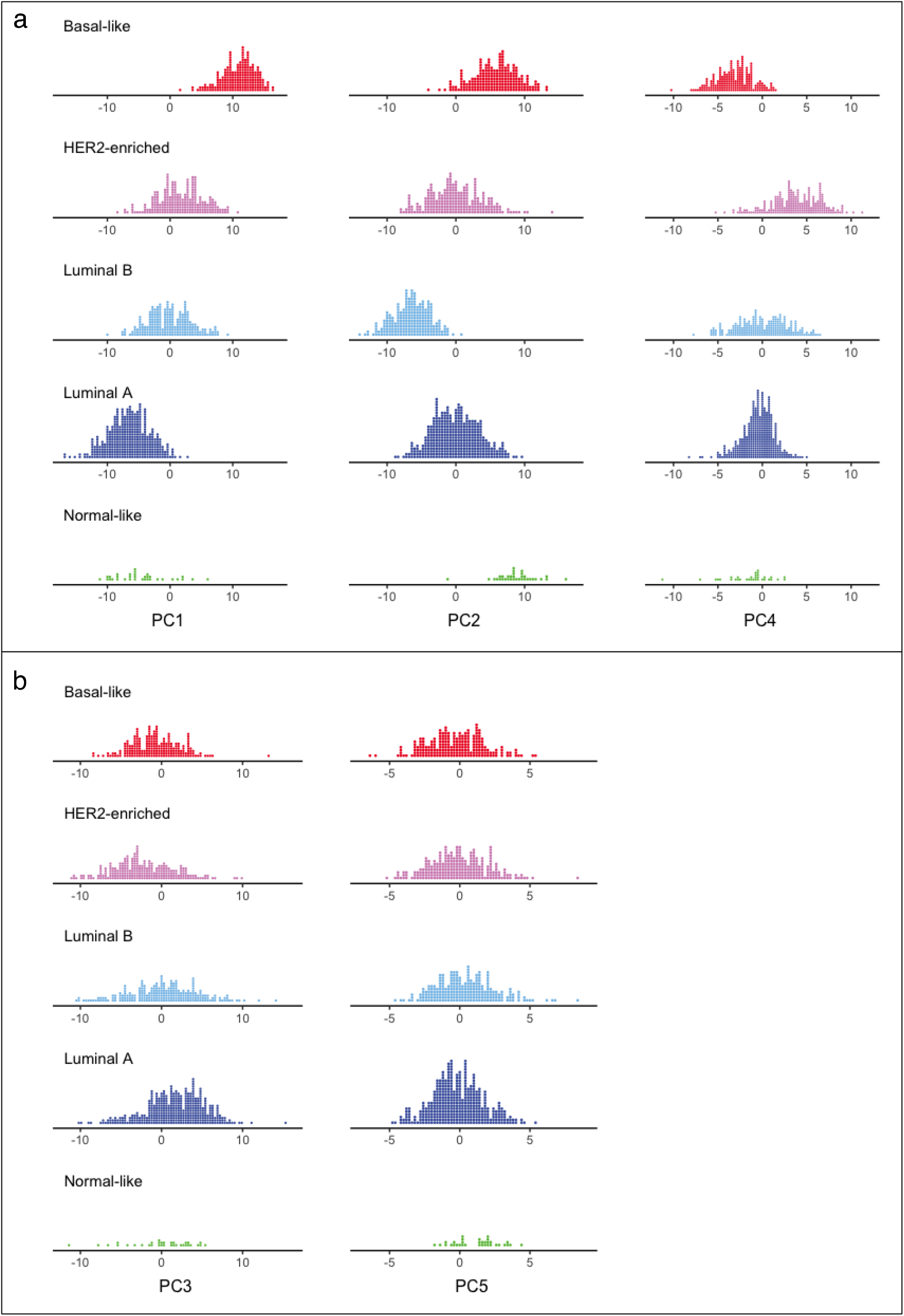
Distribution of PC scores by intrinsic subtype in LACE/Pathways. a. PC1, PC2 and PC4 capture key features of intrinsic subtypes. PC1 illustrates proliferation and ER signaling, best differentiating Basal-like from Luminal A tumors. PC2 includes a strong signal from basal cytokeratins that clearly differentiates Basal-like from Luminal B. PC4 includes a strong signal from ERBB2 and differentiates HER2-enriched tumors. Together these 3 dimensions can recapitulate intrinsic subtype clusters. b. PC3 and PC5 are novel and are not associated with intrinsic subtype.

PC1 appears to represent ER signaling and concurrently, in the opposing direction, proliferation (major coefficients for PC1 include *PGR, ESR1, NAT1, FOXA1*, Kendall’s tau with proliferation: 0.65, S1 Figure). Based on the stacked histograms (Figure 2a), PC1 clearly delineates Basal-like from Luminal A tumors. PC2 includes strong coefficients for cytokeratins *KRT5, KRT14* and *KRT17*, as well as other basal markers (*SFRP1* and *MIA*) and differentiates Basal-like from Luminal B tumors (Figure 2a). PC4 is the only component with a large coefficient for ERBB2, and contains substantial coefficients for growth factor genes (*EGFR, FGFR4* and *GRB7*). As expected, PC4 correlates well with the ERBB2 score (Kendall’s tau: 0.55, S2 Figure), and differentiates HER2-enriched tumors (Figure 2a). Together, PC1, PC2 and PC4 successfully recapitulate the 4 most common intrinsic subtypes (Figure 3).

**Figure 3.**
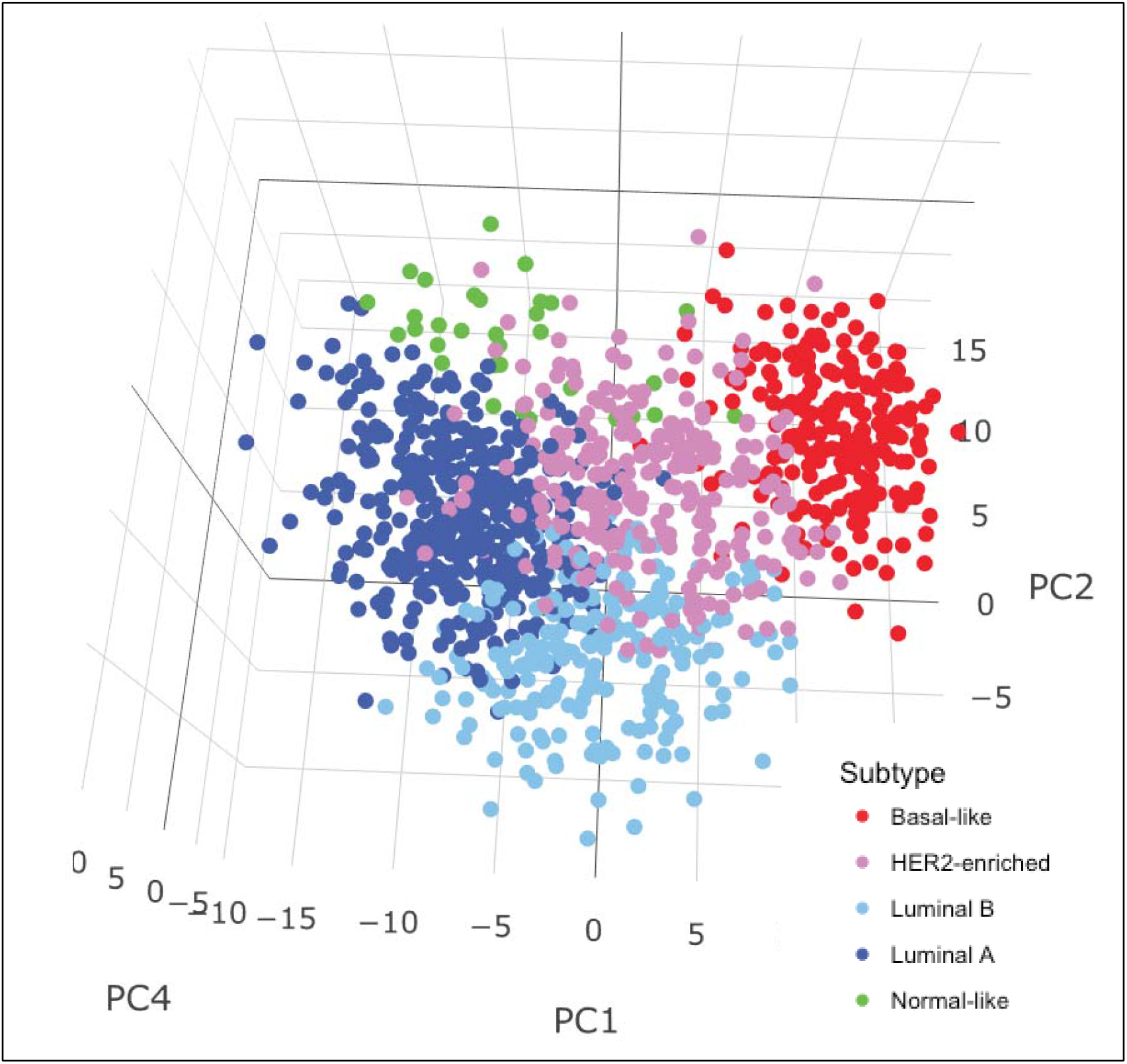
PC1, PC2, and PC4 combined recapitulate PAM50 intrinsic subtypes.

Dimensions PC3 and PC5 were novel. They were not associated with intrinsic subtypes (Figure 2b), nor were they highly correlated with PAM50 proliferation, PGR, ESR or ERBB2 scores. A notable characteristic for PC3 was that gene coefficients for basal cytokeratins were strong, but co-expressed with ER-regulated genes (atypical for any intrinsic subtype). PC5 also exhibited coefficients for *KRT17* and ER-regulated gene expression in the same direction, but otherwise was most similar to PC4, including appreciable coefficients for *EGFR, FGFR4*, and *GRB7*.

Despite extensive differences in expression technology, all PCs replicated in the TCGA RNA sequencing data (S3 Figure, S4 Figure), confirming the PC dimensions are robust and do not suffer unduly from over-fitting. In particular, the TCGA data replicated the novel dimensions PC3 and PC5. To explore whether these novel dimensions were acting as proxies for other genes, correlations between the TCGA PC3 and PC5 scores and gene expression for each of the other genes in the transcriptome were evaluated. The distribution of these correlations was Gaussian with no outliers for both PC3 and PC5. The three most highly correlated genes with PC3 score were *KRT14* (*ρ*=0.57), *KRT17* (*ρ* =0.57) and *KRT5* (*ρ* =0.55), reflecting PAM50 genes which were among the highest-ranked coefficients in the PC3 eigenvector. The most highly correlated PAM50 gene with PC5 score was *MMP11* (*ρ* =0.76) which is the highest rank coefficient in the PC5 eigenvector. Given no individual genes beyond those in the PAM50 correlated highly with either PC3 or PC5, these novel dimensions are more likely representative of a complex/pathway of multiple aberrantly expressed genes.

### Enrichment of tumor dimensions PC3 and PC5 in high-risk pedigrees

Pedigrees were not homogeneous by intrinsic subtype (Table 1, Figure 1b), nor were pedigrees significantly increased for any subtype after controlling for multiple testing. However, we did detect highly significant differences between the high-risk pedigree and the population tumors in dimensions PC3 (p<1×10^−12^) and PC5 (p=7.6×10^−8^). Furthermore, 6 pedigrees were individually significantly different from the general population for either PC3 and/or PC5 (Table 2). The significant elevation of PC3 and PC5 scores in high-risk pedigrees makes these excellent phenotypes to investigate for shared inherited susceptibility.

**Table 2:** Results of weighted t-test of means for PC3 and PC5. Comparison of pedigrees to the population (LACE/Pathways). Bonferroni corrected.

| Pedigree | n | PC3 |  | PC5 |  |
| --- | --- | --- | --- | --- | --- |
|  |  | p | t | p | t |
| 1800 | 20 | ns | 1.74 | ns | 1.609 |
| 1801 | 17 | ns | 2.03 | ns | 2.311 |
| 1808 | 24 | 0.0008 | 4.76 | 0.0129 | 3.674 |
| 1809 | 15 | 0.0778 | 3.13 | ns | 2.468 |
| 1812 | 17 | ns | 2.12 | ns | 0.305 |
| 1817 | 35 | 0.0006 | 4.55 | 0.0133 | 3.508 |
| 1818 | 20 | 0.0147 | 3.72 | ns | 0.230 |
| 1819 | 26 | 0.0001 | 5.75 | ns | 1.437 |
| 1820 | 20 | ns | 1.60 | 0.0033 | 4.352 |
| 1821 | 18 | 0.1014 | 2.92 | ns | 1.735 |
| 1822 | 31 | 0.00004 | 5.54 | ns | 0.184 |

### Shared Genomic Segment (SGS) analysis

As a pedigree-specific set, breast tumors in pedigree 1817 had significantly higher values for dimensions PC3 (p=6×10^−4^) and PC5 (p=0.013) than the population tumors (Table 2). Hence, 1817 was selected as a pedigree case-study to explore the utility of PC3 and PC5 for gene mapping. A total set of 15 women had tumors that were extreme for either PC3, PC5, or both (above the 90^th^ population percentile for the dimension). Germline DNA was available for all 15 women, either from peripheral blood or from GU breast tissue, and were germline SNP genotyped: 4 women whose tumors were extreme for both PC3 and PC5; 6 women whose tumors were extreme for only PC3; and 5 women whose tumors were extreme for only PC5. After quality control, 571,489 SNPs in 14 women were available for SGS analysis. The nine PC3-extreme breast cancer cases with SNP data were separated by 36 meioses, and the nine PC5-extreme breast cancer cases were separated by 43 meioses (Figure 1b). Ordered subsets for both PC dimensions were analyzed across the genome. Genomewide significance thresholds of 5.0×10^−7^ and 5.9×10^−7^, were established for PC3 and PC5 respectively.

One genomewide significant 0.5 Mb region (70.3–70.8 Mb, hg18) was identified for the PC3-extreme tumors at chromosome 12q15 (p=2.6×10^−8^, LOD equivalent=6.4). No genomewide significant regions were found for the PC5-extreme analysis. The PC3 12q15 locus was shared by the 8 women with the most extreme PC3 tumors (all above the 95^th^ percentile) and was inherited through 32 meioses. Only three genes reside in the SGS region: *CNOT2* (CCR4-NOT Transcription Complex Subunit 2), *KCNMB4* (Potassium Calcium-Activated Channel Subfamily M Regulatory Beta Subunit 4), and part of *MYRFL* (Myelin Regulatory Factor-Like). Notably, the gene *CNOT2* is a subunit of the CCR4-NOT complex, a global transcriptional regulator(23,24) involved in cell growth and survival(25,26). Post-hoc inspection of the PC3-extreme tumor breast cancer cases sharing the 12q15 region did not reveal any previously suggested characteristics that alternatively could have been used to identify this subset. Cases did not cluster in particular branches of the pedigree (Figure 1b), are not homogeneous for intrinsic subtype (as previously noted), and do not share similar ages at diagnosis.

### Discussion

The PAM50 gene expression assay was designed for molecular subtyping breast tumors into categorical intrinsic subtypes(12). We revisited use of these 50 gene features, defined an alternate parameterization via principal components, and identified 5 PC expression dimensions. Two of 5 expression dimensions within the PAM50 genes (PC3 and PC5) are previously unrecognized tumor characteristics and are independent of intrinsic subtypes. Key genes driving these new dimensions are also those important in architypal intrinsic types, but with atypical co-expression. Novel dimension PC3 includes substantial coefficients for ER-regulated genes and basal cytokeratin genes (*KRT5, KRT14* and *KRT17*) acting in the same direction. Luminal breast cancers are uniformly ER positive, but express cytokeratins 8 and 18(27). Whereas, Basal-like tumors are ER negative and express cytokeratins 5, 14, 17(27,28). Hence, PC3-extreme (high) tumors are a mix: luminal tumors that also express basal cytokeratins. We note myoepithelial cells comprising “normal” ducts also express basal cytokeratins, so this signature is also evident in the Normal-like subtype (Figure 2a). Proliferative tumors do not usually exhibit normal stroma contamination and basal cytokeratins are therefore an uncharacteristic feature in Luminal B and HER2-enriched tumors, which were included in the PC3-extreme tumors. This suggests the basal cytokeratin expression comes from the ER positive tumor epithelial cells, rather than normal stroma contamination; however, further investigation is necessary to confirm this atypical luminal tumor expression of basal cytokeratins.

While comparisons of high-risk pedigree and population tumors did not support a ‘same-gene-same-intrinsic-subtype’ hypothesis, a ‘same-gene-same-tumor-dimension’ hypothesis was supported. Although not explicitly shown here, it is highly likely that PC, which clearly differentiates Basal-like tumors in its extreme tail (S1 Figure) would be significantly enriched in *BRCA1* pedigrees. Here, novel breast tumor dimensions, PC3 and PC5, were found to be significantly enriched in *non-BRCA1/2* high-risk pedigrees. This provides potential new heritable breast cancer phenotypes and new opportunities to identify genetic susceptibility loci through reduced heterogeneity and increased statistical power. Consistent with this hypothesis, we presented a proof-of-concept gene-mapping case-study using PC3-extreme tumors in one high-risk pedigree which identified a genomewide significant 0.5 Mb region at 12q15 that inherited to the 8 most PC3-extreme breast cancer cases across 32 meioses (p=2.8×10^−8^). Of the three genes residing in the 0.5 Mb region, *CNOT2* is a compelling candidate because of its role as a regulatory protein of the CCR4 (carbon catabolite repressor-4)-NOT (negative on TATA) complex, which functions as a master regulator of transcription, translation and mRNA stability(29,30). CCR4-NOT and *CNOT2* have been demonstrated to function in the regulation of DNA damage response, cell cycle progression, DNA replication stress response, and control of cell viability(26,31–33). A transcriptional module of *CNOT2* has also been correlated with heritable susceptibility of metastatic progression in a mouse model of breast cancer. Previously direct involvement of *CNOT2* in metastasis has been demonstrated; knockdown of *CNOT2* enhanced and overexpression of *CNOT2* attenuated lung metastasis of mouse mammary tumor cells(25). An attractive possibility is that a germline risk modifies *CNOT2* expression, leading to dysregulation of mechanisms controlling cell growth and DNA damage repair. The natural next step will be to determine specific genetic variants in our 12q15 locus and assess effect on *CNOT2*.

Beyond our application in gene-mapping, a PC dimension approach to reparameterize expression of pre-selected gene features may have utility in other domains. Principal components are orthogonal measures, providing independent variables for multi-variate modeling. This flexibility has the potential for increased power over a single variable categorical approach by allowing multiple expression dimensions to be modeled simultaneously. In particular, other study designs using the PAM50 expression for tumor characterization can immediately explore the PCs described here proving additional opportunities to identify novel clinical or therapeutic associations. Furthermore, the deeper appreciation of the gene expression dimensions of breast tumors uncovered here may be useful in illuminating functionally important tumor pathways, or particular tumor evolutions.

In summary, we have revisited interpreatation of PAM50 gene expression, identified and replicated 5 orthogonal PC dimensions, and discovered two novel breast tumor dimensions with significant evidence for underlying genetic heritability. Based on one of these novel tumor dimensions, we have mapped a genomewide significant breast cancer locus at 12q15 and present a compelling breast cancer susceptibility candidate gene, *CNOT2*. The strong statistical significance achieved by the mapping of this new breast cancer susceptibility locus harkens back to the era of pedigree gene-mapping successes. These novel tumor dimensions may, indeed, reduce germline genetic heterogeneity and hold promise for a new wave of susceptibility gene discovery in breast cancer. Furthermore, the appreciation of all five expression dimensions from the PAM50 assay lends additional informative variables that can be assessed immediately, with potential for new discoveries in other study designs where molecular phenotypes are important, such as, treatment response and clinical outcome studies.

### Data availability

Pedigree expression data referenced in this study will be available in the Gene Expression Omnibus database. TCGA expression data is available from the NCA Genomic Data Commons Data Portal (Project ID: TCGA-BRCA).

### Computer code

The software used for SGS analysis is available at https://gitlab.com/camplab/sgs and https://gitlab.com/camplab/jpsgcs.

### Funding

This work was supported by the National Cancer Institute [grant numbers CA163353, CA129059, CA105274, CA195565].

## Acknowledgements

This work was aided by the Genomics Core Facility and through the computational resources and staff expertise provided by the Center for High Performance Computing at the University of Utah. We thank the Pedigree and Population Resource of the Huntsman Cancer Institute, University of Utah (funded in part by the Huntsman Cancer Foundation) for its role in the ongoing collection, maintenance and support of the UPDB. Finally, we thank the participants and their families who make this research possible.

## Supplementary Information

**S1 Figure.**
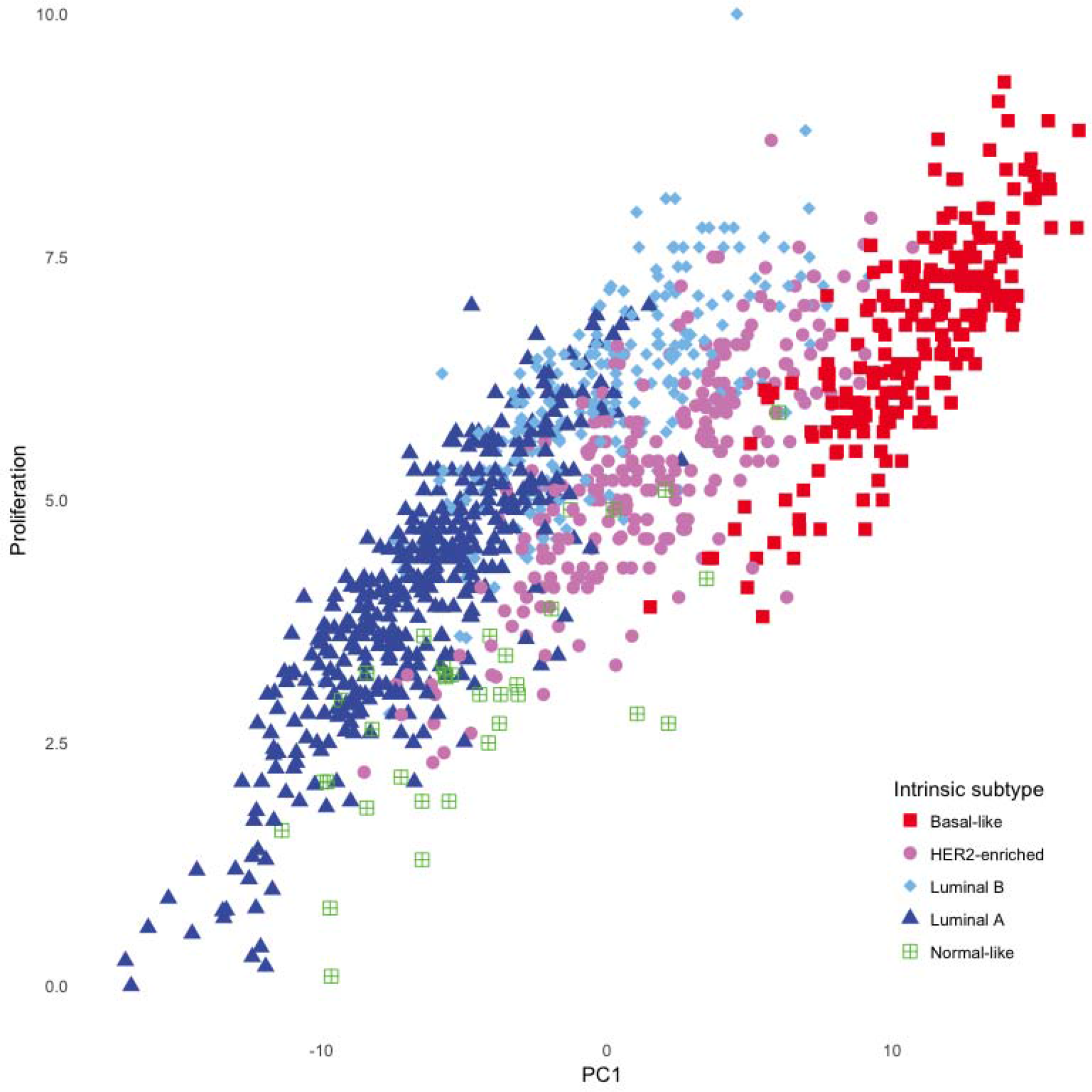
Correlation of PAM50 proliferation score with PC1 in LACE/Pathways.

**S2 Figure.**
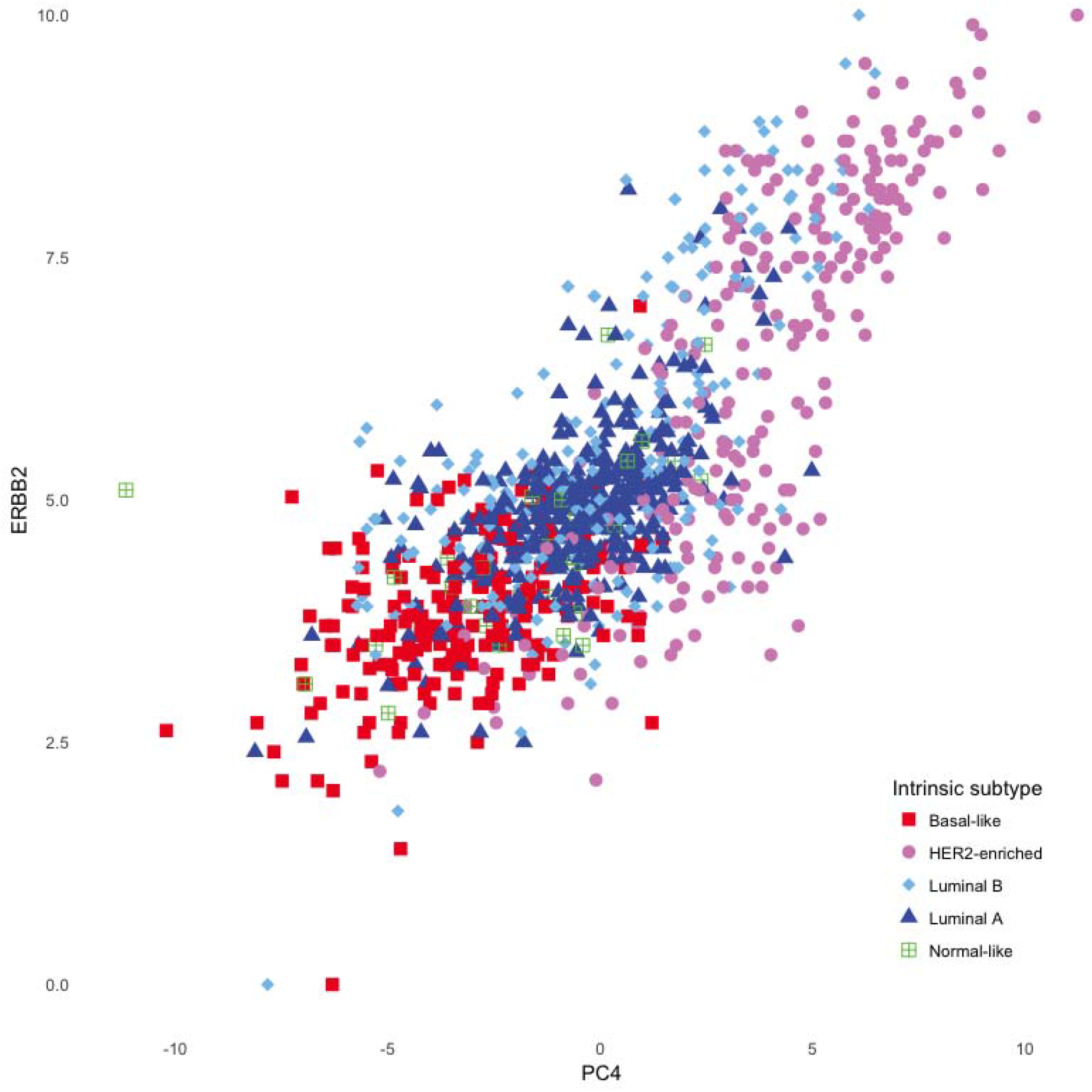
Correlation of PAM50 ERBB2 score with PC4 in LACE/Pathways.

**S3 Figure.**
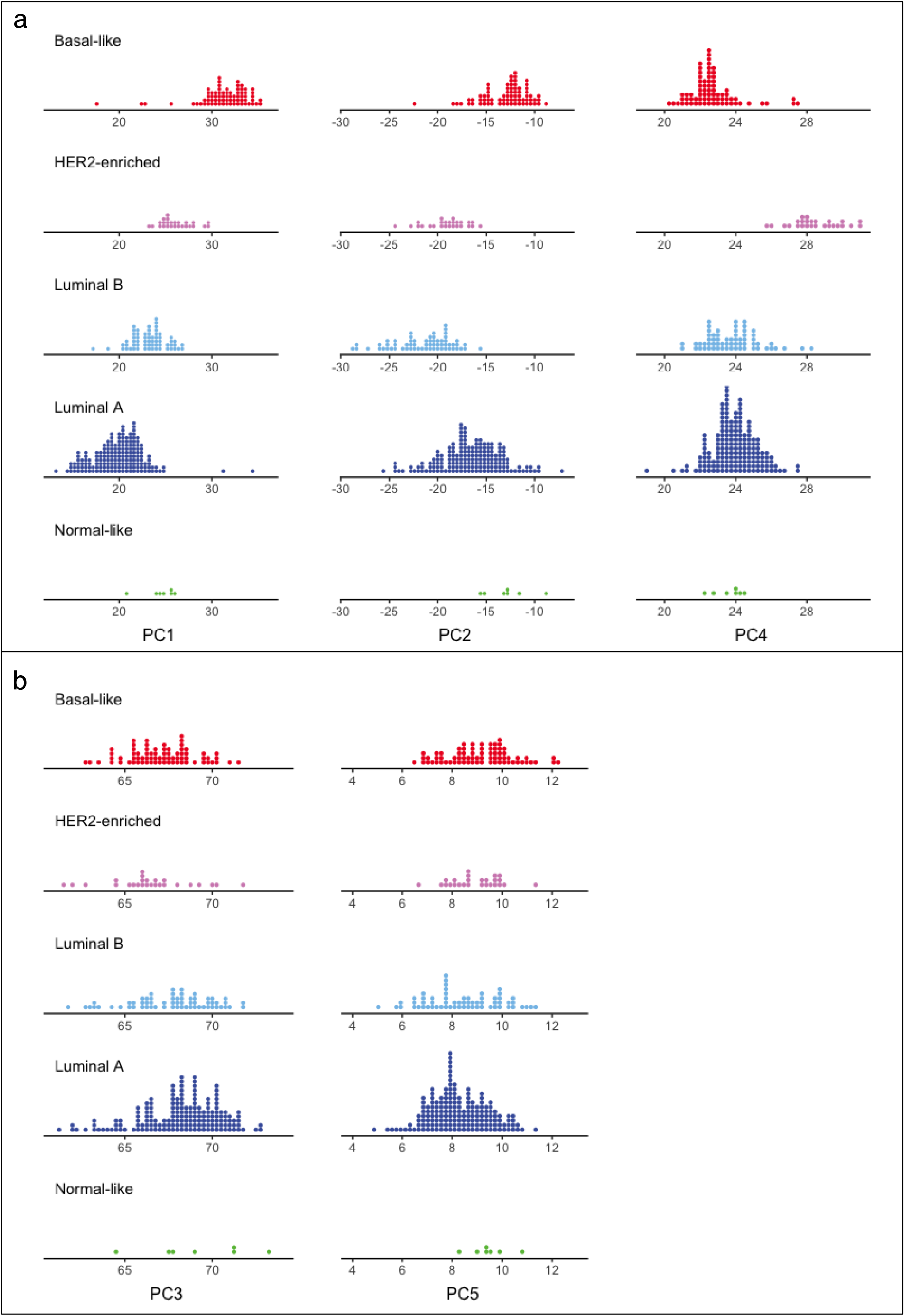
Distribution of PC scores by intrinsic subtype replicate in TCGA expression data. **a.** PC1, PC2 and PC4 capture key features of intrinsic subtypes. **b.** PC3 and PC5 are independent of intrinsic subtype.

**S4 Figure.**
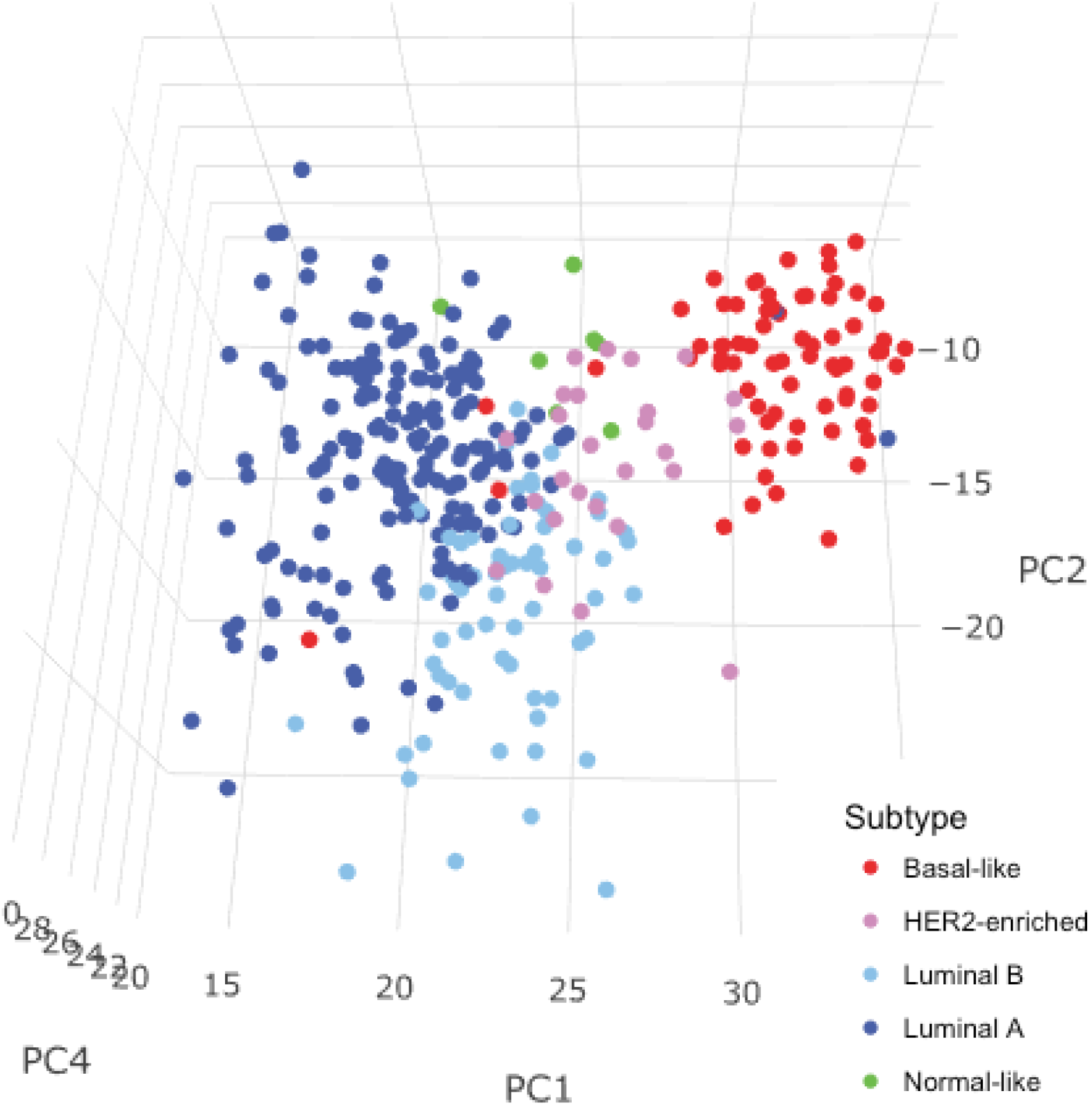
Intrinsic subtypes remain evident in TCGA breast tumors based on direct application of the PC1, PC2, and PC4 equations (linear combinations of expression) derived from PAM50 (LACE/Pathways) to RNAseq data from TCGA breast tumors.

**S1 Table.** Eigenvectors and eigenvalues of principal components 1-5.

